# Leveraging the genetic diversity of trout in the rivers of the British Isles and northern France to understand the movements of sea trout (*Salmo trutta* L.) around the English Channel

**DOI:** 10.1101/2024.05.10.593494

**Authors:** R. Andrew King, Charlie D. Ellis, Dorte Bekkevold, Dennis Ensing, Thomas Lecointre, Daniel R. Osmond, Adam Piper, Dylan E. Roberts, Sophie Launey, Jamie R. Stevens

## Abstract

Populations of anadromous brown trout, also known as sea trout, have suffered recent marked declines in abundance due to multiple factors, including climate change and human activities. While much is known about their freshwater phase, less is known about the species’ marine feeding migrations. This situation is hindering the effective management and conservation of anadromous trout in the marine environment. Using a panel of 95 single nucleotide polymorphism markers we developed a genetic baseline, which demonstrated strong regional structuring of genetic diversity in trout populations around the English Channel and adjacent waters. Extensive baseline testing showed this structuring allowed high-confidence assignment of known-origin individuals to region of origin. This study presents new data on the movements of anadromous trout in the English Channel and southern North Sea. Assignment of anadromous trout sampled from 12 marine and estuarine localities highlighted contrasting results for these areas. The majority of these fisheries are composed predominately of stocks local to the sampling location. However, there were multiple cases of long-distance movements of anadromous trout, with several individuals originating from rivers in northeast England being caught in the English Channel and southern North Sea, in some cases more than 1000 km from their natal region. These results have implications for the management of sea trout in inshore waters around the English Channel and southern North Sea.

## 1 Introduction

Brown trout (*Salmo trutta* L.) is a ubiquitous fish species found naturally over much of Europe, north Africa and western Asia in a wide range of river types (Kershner, Williams, Gresswell, & Lobón-Cerviá, 2019). Across this range, brown trout show a great range of morphologies (Ferguson & Prodöhl, 2022; Verspoor, Coulson, Greer, & Knox, 2019) and genetic variants (Bernatchez, 2001; Ferguson, 1989; Ferguson & Taggart, 1991; King, Hillman, Elsmere, Stockley, & Stevens, 2016; Quéméré et al., 2016; Vilas, Bouza, Castro, López, & Martínez, 2010). These genetic variants can often be highly localised, with distinct patterns of genetic variation between fish inhabiting different parts of a catchment and/or adjacent rivers (Bekkevold et al., 2020; Bouza, Arias, Castro, Sánchez, & Martínez, 1999; Ferguson, 1989; Griffiths, Koizumi, Bright, & Stevens, 2009; King, Stockley, & Stevens, 2020). These levels of significant genetic separation allow the recognition of distinct populations and reflect both the phylogeographic history of the species (Bernatchez, 2001; Cortey, Vera, Pla, & García-Marín, 2009; McKeown, Hynes, Duguid, Ferguson, & Prodöhl, 2010) and more recent events that have acted to restrict or eliminate gene flow, e.g., the construction of dams and weirs (King et al., 2020; Osmond, King, Russo, Bruford, & Stevens, in press), leading to the emergence of distinct genetic signatures due to drift and adaptation. In turn, these distinct populations can be used as OTUs for the assessment of straying (King et al., 2016) in anadromous individuals (hereafter referred to as sea trout) and for tracing the at-sea movements of fish (Bekkevold et al., 2021; Koljonen, Gross, & Koskiniemi, 2014; Prodöhl et al., 2017). Both are achieved by assigning sea trout back to their population or region of origin based on similarities between the genotypes of the migratory form (sea trout) and the population genetic signature of resident trout in different candidate rivers/regions of origin.

The English Channel is one of the busiest waterways in Europe for both commercial and recreational fishing, cross-Channel trade and as a navigation route from the Atlantic to the southern North Sea and the Baltic (Glegg, Jefferson, & Fletcher, 2015). Along its length several major rivers flow into it, including the Seine and, historically, it forms the route of the paleo-Channel River (Lericolais, Auffret, & Bourillet, 2003). Thus, many of the rivers of this region have a common history, beginning as tributaries of the much larger ancient Channel River and sharing riverine geologies. Similarly, the trout of this region have a shared history dating from before the last glacial maximum (Bernatchez, 2001; McKeown et al., 2010) and have been affected by rising sea levels after the LGM, leading to the separation of many former Channel River tributaries into distinct catchments.

More recently, populations of both trout and Atlantic salmon have been severely affected by human-related activities, including targeted estuarine net fisheries, changes to river navigability and barriers to upstream movement (weirs, dams), point-source and diffuse pollution, loss of spawning habitat and many stocking and translocation events (Losee et al., 2024; Nevoux et al., 2019). It is this combination of historic and contemporary factors that have shaped the present mosaic of genetic groupings of trout in rivers on both sides of the English Channel and in the southern North Sea (King et al., 2016; King et al., 2020; Quéméré et al., 2016). Research has been able to inform on the impact of many of the factors driving population level variation in trout, particularly those acting in the freshwater phase of the trout lifecycle (King et al., 2020; Paris, King, & Stevens, 2015). However, trout – unlike salmon– exhibit a continuum of life history variation from fully resident through freshwater migration to fully anadromous individuals (Ferguson, Reed, Cross, McGinnity, & Prodöhl, 2019).

There is a long history of studies investigating the marine distribution of different stocks and the mixed-stock nature of marine fisheries in anadromous salmonids at different spatial scales (Cormack & Skalski, 1992; Tucker et al., 2009). Recently, there has been extensive investigation of the marine distribution of different Atlantic salmon stocks and the mixed stock nature of targeted marine fisheries assessed using genetic baselines (Bradbury et al., 2015; Gilbey et al., 2021; Gilbey et al., 2017); to date, however, there have been only a limited number of similar studies on sea trout (Bekkevold et al., 2021; Koljonen et al., 2014; Prodöhl et al., 2017). Unlike Atlantic salmon, however, anadromous trout are thought to feed more locally to their natal rivers (Jonsson & Jonsson, 2014; Malcolm, Godfrey, & Youngson, 2010; Potter, Campbell, Sumner, & Marshall, 2017), rather than migrating long distances to offshore feeding grounds in the north Atlantic (Gilbey et al., 2021; Gilbey et al., 2017). Nonetheless, several tagging and tracking studies have reported highly variable degrees of movement, including longer migrations of limited numbers of individuals (Hawley, Urke, Kristensen, & Haugen, 2024; Kallio-Nyberg, Saura, & Ahlfors, 2002; Malcolm et al., 2010; Potter et al., 2017). Additionally, distinct regional differences in migration patterns have been reported (Potter et al., 2017).

With anadromous salmonids being subject to multiple stressors, both in their freshwater and marine environments, many species have suffered marked declines in abundance over recent decades (ICES, 2013). While management and conservation measures for trout in freshwater, including knowledge of when and where to implement such measures, are now relatively well understood, an understanding of how, when and where to implement protection measures for trout in the marine environment is much less advanced. Similar to Atlantic salmon (Gillson et al., 2022), within the marine environment, stressors of sea trout include aquaculture, coastal developments (i.e. tidal lagoons, inshore and offshore wind farms), and by-catch in non-target fisheries (Nevoux et al., 2019; Thorstad et al., 2016).

Given the importance of anadromous individuals to the resilience of trout populations (Goodwin, King, Jones, Ibbotson, & Stevens, 2016), effective conservation and management of such populations requires extensive information on species biology, behaviour, life cycle and the challenges they face at different life history stages (Nevoux et al., 2019; Whelan et al., 2017), including knowledge of when and where sea trout go during their marine migrations (O’Sullivan et al., 2022; Thorstad et al., 2016). Of particular relevance is the incidence of individuals taken as by-catch in non-target marine fisheries; again, data on this specific to sea trout are very poor (Elliott et al., 2023).

In this study, we constructed a genetic baseline for trout sampled from 107 rivers around the English Channel, southern Irish Sea and southern North Sea based on 95 single nucleotide polymorphism (SNP) markers. Our objectives were (1) to catalogue the structuring of, and genetic variation between, trout populations in these areas, (2) to assess the scale at which reliable assignment to the baseline could be achieved using leave-one-out analyses and genotypes from known-origin individuals, and (3) to investigate the stock composition of sea trout sampled from multiple marine and estuarine locations along the English Channel, Bristol Channel and southern North Sea coasts of England, France and the Netherlands.

## 2 Materials & Methods

### 2.1 Sample collection

For baseline construction, adipose finclip or scale samples from juvenile resident trout were obtained from various sources (Supplementary Table 1). Samples were collected during routine electrofishing surveys in the UK and Ireland by the Environment Agency in England, Inland Fisheries Ireland or, specifically, by the SAMARCH project team (www.samarch.org) and in France by INRAE U3E Unit, Office Français de la Biodiversité, Bretagne Grands Migrateurs, Seinormigr and Fédération Départmentale de Pêche et de Protection du Milieu Aquatique 14, 22, 27, 29, 35, 50, 62, 76 and 80 as part of inventory surveys. Samples from two Danish rivers consisted of mature adults collected on spawning sites by a team from the Technical University of Denmark - details in Bekkevold et al. (2020).

Scale and finclip samples from 398 sea trout were obtained from commercial and recreational fisheries from English, French and Dutch coastal and estuarine areas (Supplementary Materials and Supplementary Figure 1). These collections represent a range of samples caught in targeted commercial salmonid netting activities (*i.e.* TT and EAN), as by-catch in commercial fisheries targeting non-salmonids (*i.e.* RYE), recreational fisheries (*i.e.* OUS and MER) or targeted sampling (*i.e.* KIM and COR) undertaken specifically for the SAMARCH research project (www.samarch.org). Details of these fisheries are given in the Supplementary Materials.

### 2.2 Molecular Methods

Genomic DNA was extracted using the HotSHOT method of Truett et al. (2000) for southern UK and Irish samples, Omega Biotek E.Z.N.A. kits for NE English and Danish samples and NucleoSpin^®^ 96 Tissue kits (Macherey-Nagel) for French samples. All individuals were genotyped at 95 biallelic single nucleotide polymorphism (SNP) loci (Osmond, King, Stockley, Launey, & Stevens, 2023) on the Fluidigm EP1 Genotyping System using 96.96 Dynamic Genotyping Arrays and scored using the Fluidigm SNP Genotyping analysis software. Genotype plots of each locus were manually inspected for quality of individual genotyping and clustering. Individual points that fell outside of the heterozygote or homozygote genotype clusters were considered to have poor quality data and left uncalled for that locus (Clemento, Abadia-Cardoso, Starks, & Garza, 2011). Individual genotypes with more than 5 uncalled loci were excluded from subsequent analyses. Each run included two positive (individuals of known genotype) and two negative (no DNA) controls.

### 2.3 Data Quality Assurance

Juvenile salmonid populations can sometimes be characterised by large numbers of closely related individuals, *i.e.* full-sibs (Goodwin et al., 2016), the presence of which can lead to biases in the inference of population structure (Anderson & Dunham, 2008) and genetic stock identification (Östergren, Palm, Gilbey, & Dannewitz, 2020). To assign sibship within each sample of fish we used a maximum-likelihood method, implemented in COLONY v 2.0 (Jones & Wang, 2010). Settings were: high precision medium length run, assuming both male and female polygamy without inbreeding and a conservative 0.5% error rate for both scoring error rate and allelic dropout rate. To check for consistency, analyses were run twice using different random number seeds. Full sibs were trimmed from the data set using Waples and Anderson’s (2017) Yank-2 method – all but two random members of families with three or more individuals were removed.

Linkage disequilibrium (LD) between all pairs of loci within each population was tested using GENEPOP v3.4 (Raymond & Rousset, 1995). Significance was estimated using a Markov chain method using default parameters (1000 de-memorisations, 100 batches and 1000 iterations). False Discovery Rate (FDR, (Benjamini & Hochberg, 1995)) was used to correct significance levels for multiple comparisons - https://www.multipletesting.com, Menyhart, Weltz & Gyorffy (2021). Using GenoDive v3.03 (Meirmans, 2020), deviations from Hardy-Weinberg Equilibrium (HWE) for each locus and population was assessed using Nei’s (1987) heterozygosity-based G_IS_ estimator with significance based on 999 permutations.

### 2.4 Basic measures of genetic diversity

GenoDive v3.03 (Meirmans, 2020) was used to calculate observed (H_O_) and unbiased expected heterozygosity (H_E_) and Weir & Cockerman’s (1984) estimator of F_ST_ were calculated with significance of F_ST_ values determined using 999 bootstrap replicates.

### 2.5 Population Genetic Structure and Identification of Reporting Groups

Depending on location, salmonid fisheries often target mixed stocks of fish with ‘stocks’ comprising multiple, geographically proximate and genetically similar rivers (Moran & Anderson 2019). To investigate population genetic structuring of trout populations, we performed two analyses. Firstly, we used STRUCTURE v 2.3.4 (Pritchard, Stephens, & Donnelly, 2000) which implements a Bayesian-based Markov Chain Monte Carlo (MCMC) model-based clustering method to jointly delineate *K*, the number of partitions of the data set and q, the proportion of each individual’s genome originating from each of the *K* partitions. STRUCTURE was run with a burn-in of 100 000 iterations followed by 250 000 iterations with the number of inferred populations (*K*) ranging from one to 15. Ten independent runs were performed using the admixture model with correlated allele frequencies and not using the population of origin information as a prior. We used the Δ*K* method of Evanno *et al*. (2005) to determine the most likely number of clusters. Hierarchical analyses were performed, based on the Δ*K* results for the full data set, to identify finer-levels of structure. Where the number of rivers in a hierarchical analysis was less than 15, the maximum *K* was set at N_rivers_ + 1. POPHELPER v1.0.6 (Francis, 2017) was used to calculate Δ*K* and to visualise the consensus data after alignment of multiple runs at optimum *K* values using CLUMPP v 1.1.2 (Jakobsson & Rosenberg, 2007).

A neighbour-joining dendrogram based on Cavalli-Sforza and Edwards (1967) chord distance (D_CE_) was used to identify population-level genetic structure. The dendrogram was constructed and visualised using POPULATIONS v1.2.32 (Langella, 1999) and MEGA v6 (Tamura, Stecher, Peterson, Filipski, & Kumar, 2013), respectively. Baseline reporting groups, upon which subsequent assignments would be based, were identified using a combination of the STRUCTURE and neighbour-joining analyses.

### 2.6 Genetic Stock Identification Analyses

We employed two widely utilised pieces of assignment software for the mixed stock analyses (MSA) and individual assignment (IA) of sea trout caught in estuarine and marine waters to both individual river and reporting groups as defined in the population structure analyses (see Results section). cBayes (Neaves, Wallace, Candy, & Beacham, 2005) implements the Bayesian procedures of Pella & Masuda (2001). For stock composition estimation, eight 50 000-iteration Markov Chain Monte Carlo (MCMC) chains were run, with initial values set at 0.9 for each chain for different samples. Means and 95% confidence intervals of the estimated stock contributions were determined from the combined final 1000 iterations from each chain. *RUBIAS* uses a Bayesian conditional genetic stock identification model to provide mixture proportion estimates and assign individuals to population/stock of origin (Moran & Anderson, 2019). Assignment proportions and their 95% credible intervals were generated using the MCMC method based on 100 000 sweeps following a burn-in of 10 000 sweeps.

We used two tests to assess the accuracy of assignments to our SNP baseline. Firstly, Leave-One-Out (LOO) analysis, as implemented in *RUBIAS*, was used to assess assignment accuracy and efficiency. Secondly, we assessed the mixed-stock and individual assignment of 436 individuals of known origin from 25 baseline rivers using both cBayes and *RUBIAS*. Full details of these tests and their results are given in Supplementary Materials.

Mixed stock analysis and individual assignment to reporting group for the 12 marine and estuarine derived collections of sea trout were estimated using both cBayes and *RUBIAS*. Analyses were run using the conditions given above.

Least-cost migration distances for each marine-caught sea trout were calculated using the marmap R package (Pante & Simon-Bouhet, 2013). For the East Anglian and Dutch fishery samples where fish were sampled from multiple locations, we took the approximate midpoint between the extreme sampling locations on each stretch of coastline. For regional level assignments, we calculated the minimum, maximum and average distance that fish could have migrated from a river of origin within a reporting group to the marine sampling location.

## 3 Results

### 3.1 Data quality

A total of 4085 individuals were genotyped at 95 SNP loci. Comparison of genotypes from repeated samples gave an error rate of 0.0014% (46 mismatches from 31 920 allele calls). In total, 98 individuals were removed after failing to be genotyped at ≥6 loci. The number of full-sib families per baseline sample ranged from zero to nine (mean families per river = 2.48). The maximum number of individuals in any full-sib family was ten. In total, 125 full-sib individuals were removed following analysis with the program COLONY. The final dataset comprised 3067 baseline, 436 known origin and 371 marine-/estuarine-caught sea trout.

After FDR correction, 32 pairs of loci (out of a total of 477 755 pairwise comparisons) were in significant linkage across the 107 baseline samples. There were 354 significant deviations from HWE (out of a total of 10 165 baseline sample/locus combinations). As none of these significant results showed any consistent patterns across loci or baseline samples, all loci and samples were retained for further analyses.

### 3.2 Population genetic structure

Global F_ST_ was 0.109 (p=0.001). Pairwise F_ST_ values ranged from zero (p=0.512) between the East Looe and West Looe rivers in southern Cornwall to 0.266 (p=0.001) between the Horn (Bretagne) and Sow (southeast Ireland) rivers.

The results of the STRUCTURE and neighbour-joining analyses were in broad agreement with both identifying a high degree of regional structuring within the 107 baseline rivers, with neighbouring rivers being genetically more similar to each other, sometimes over long stretches of coastline. The neighbour-joining analysis identified 13 geographical structured groups of rivers (Figure 1) with the number of rivers per group ranging from two (DENMARK) to 20 (DEVCORN). STRUCTURE identified *K*=2 (Δ*K*=122.4) as the most likely partition of the full dataset, splitting the rivers into western and eastern groupings (Figure 2). Subsequent hierarchical analyses identified further subdivision within both the western and eastern groups and broadly recovered the same population groupings as found in the Neighbour-joining analysis (Figure 2). STRUCTURE also highlighted that the distinction between genetic groups tended to be geographically limited, for example, in Britain between the Hampshire Basin and southeast English rivers (Figure 2) and in France between the rivers of Lower and Upper Normandy (Figure 2).

**FIGURE 1.**
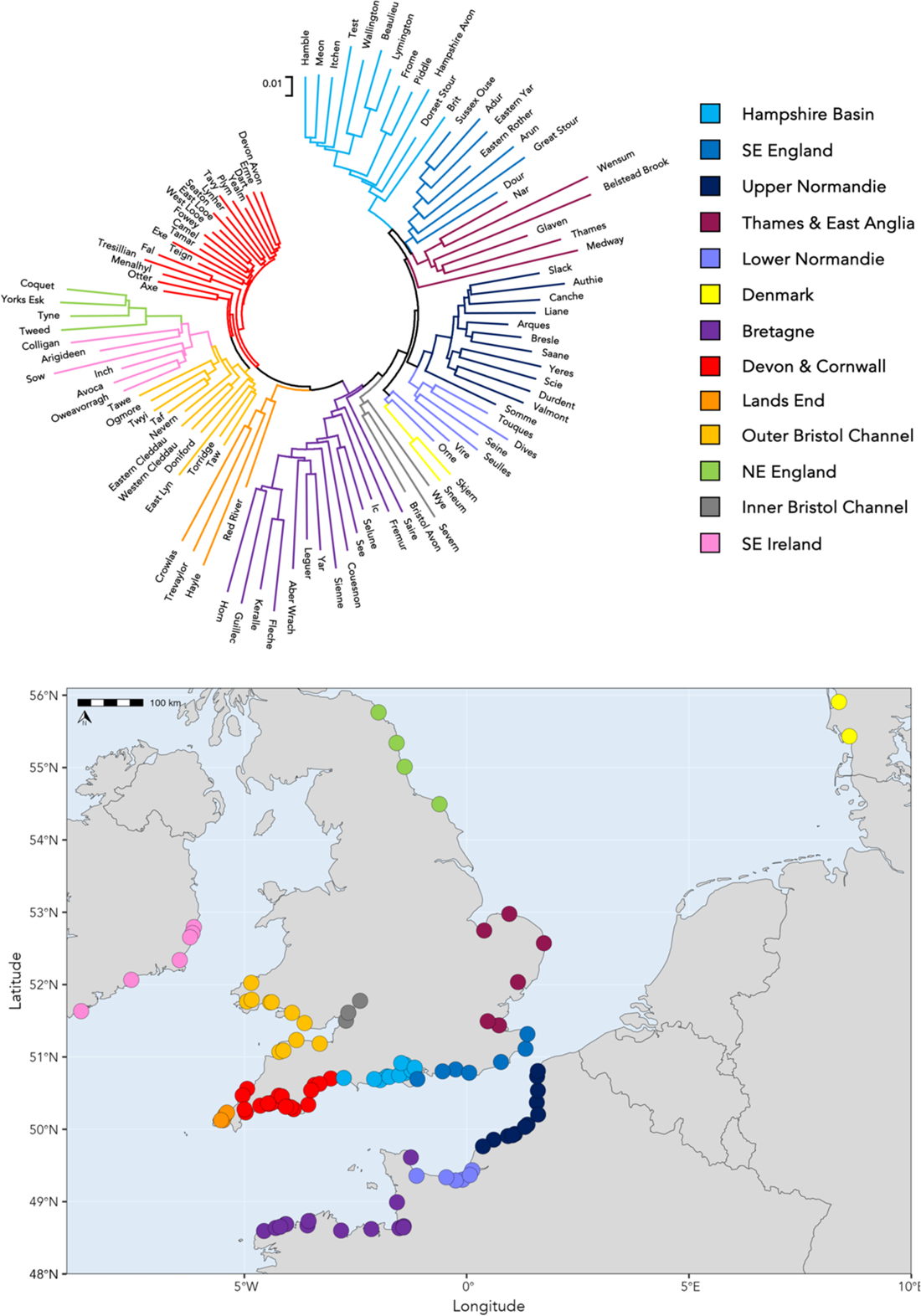
Unrooted neighbour-joining (NJ) dendrogram, based on Cavalli-Sforza and Edwards’ chord distance (D_CE_), showing relationships between the 107 resident trout populations sampled for the SNP baseline. Branches are colour coded by reporting group. The map gives the location of the mouth of each sampled river with coloured points giving reporting group membership as determined the NJ dendrogram. Full sample site details are given in Supplementary Table 1.

**FIGURE 2.**
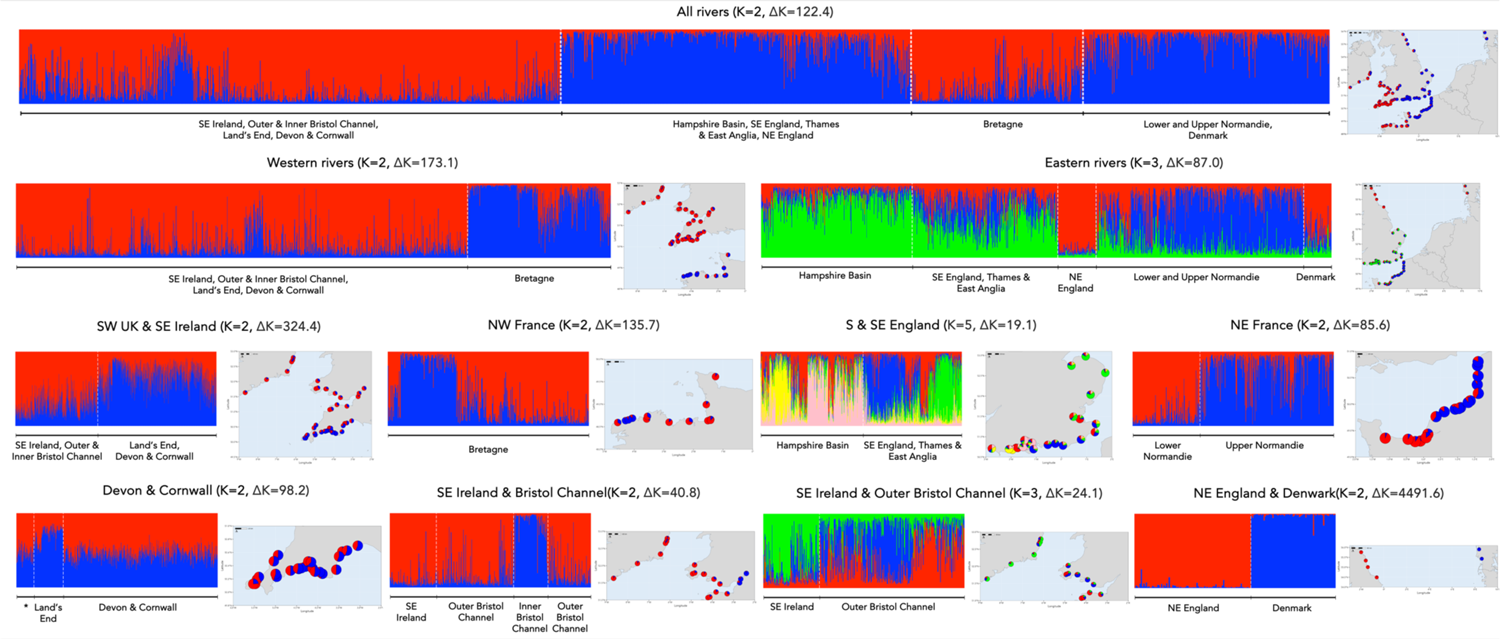
Results of the hierarchical STRUCTURE analysis for the 107 resident trout baseline rivers. Results of each STRUCTURE analysis are shown as bar plots with vertical columns represent the assignment probabilities of individuals to each of the *K* inferred clusters. For clarity, results are plotted by reporting groups rather than individual rivers. Maps show the location of each sampled river with pie charts giving the population-level assignment to each genetic cluster. Plots of Δ*K* values for each analysis are given in Supplementary Figure 5.

### 3.3 Baseline testing

Based on the regional structuring identified in the STRUCTURE and neighbour-joining analyses, we identified 13 groups of rivers (hereafter referred to as reporting groups), with the addition of a group of French hatchery populations, as the basis for the baseline testing and assignment of sea trout. Results of the initial baseline testing are given in detail in the Supplementary Materials. Briefly, LOO analysis found generally high levels (>85%) of assignment accuracy and efficiency to reporting group (Supplementary Figure 2).

Conversely, assignment success to individual rivers was highly variable. For some rivers assignment had very high (>95%) accuracy and efficiency, *i.e.* SEV, WEN, TYN (Supplementary Figure 3), however, most rivers demonstrated much lower assignment success. For example, for many of the rivers in the Devon & Cornwall reporting group (DEVCORN) accuracy and efficiency of assignment to an individual river was below 50% (Supplementary Figure 3). Mixed-stock and individual assignment of the known-origin collections showed similar trends to the LOO analysis, with collections assigning strongly to their region of origin and highly variable success of assignment to river of origin (Supplementary Figure 4, Supplementary Tables 2 and 3). There were also clear differences in the ability of *RUBIAS* and cBayes to correctly assign collections and individual fish to their rivers of origin (Supplementary Figure 4). Based on these results, here we report only regional mixed-stock and individual assignments for the 12 marine- and estuarine-caught collections determined using cBayes. However, cBayes MSA and IA results of assignment to river of origin and *RUBIAS* results for both regional and river MSA and IA are presented in Supplementary Tables 4 and 5.

### 3.4 Assignment of marine and estuarine collections

Assignment of the 12 collections of marine and estuarine sampled sea trout showed contrasting patterns of assignment. The four estuarine collections (TT, TAM, PLH & OUS, Figures 3 and 4) showed very little evidence of mixing of fish from different reporting groups, with each collection being dominated by migratory fish from the same reporting group as that to which the sampled estuaries belonged (Figures 3 and 4, Supplementary Tables 4 and 5). For example, the majority of sea trout sampled in the Taw/Torridge estuary belonged to the OUTBRCH reporting group with a single individual assigning strongly to the DEVCORN reporting group (Figures 3 and 4, Supplementary Tables 4.09 and 5.09). Likewise, 29 out of 30 fish sampled in a recreational sea trout rod fishery in the tidal reaches of the Sussex Ouse, a member of the SEENG reporting group, assigned to that reporting group. The remaining individual had strongest assignment to the NEENG reporting group (Supplementary Table 5.12).

**FIGURE 3.**
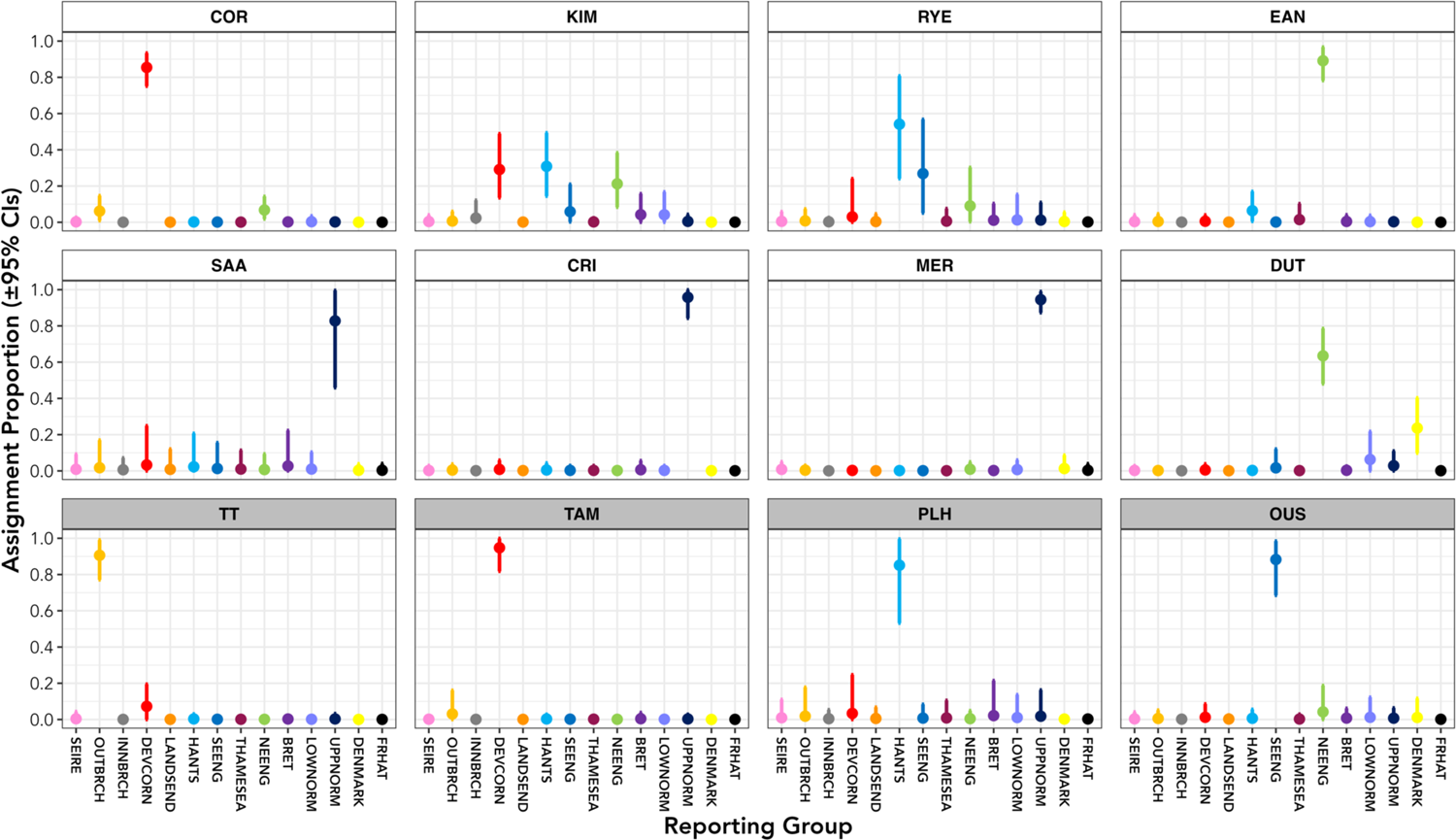
Mean estimated stock composition assigned to reporting group of origin, with 95% confidence intervals, for eight marine (white chart header) and four estuarine (grey chart header) collections of anadromous trout. Reporting regions are colour coded as given in Figure 1. Marine collection abbreviations: COR – southern Cornwall; KIM – Kimmeridge Bay; RYE – Rye Harbour; EAN – East Anglian drift-net fishery; SAA – Saâne illegal nets; CRI – Criel-sur-Mer recreational beach nets; MER – Mers-les-Bains and Le Tréport recreational beach nets; DUT – Dutch commercial fishery by-catch. Estuarine collection abbreviations: TT – Taw/Torridge shared estuary; TAM – River Tamar tidal limit fish trap; PLH – Poole Harbour; OUS – Sussex Ouse estuary recreational rod fishery. Reporting group abbreviations: SEIRE – southeast Ireland; OUTBRCH – outer Bristol Channel; INNBRCH – inner Bristol Channel; DEVCORN – Devon and Cornwall; LANDSEND – Land’s End complex; HANTS – Hampshire Basin; SEENG – southeast England; THAMESEA – River Thames and East Anglia; NEENG – northeast England; BRET – Bretagne; LOWNORM – Lower Normandy; UPPNORM – Upper Normandy; DENMARK – Denmark; FRHAT – French hatchery populations.

**FIGURE 4.**
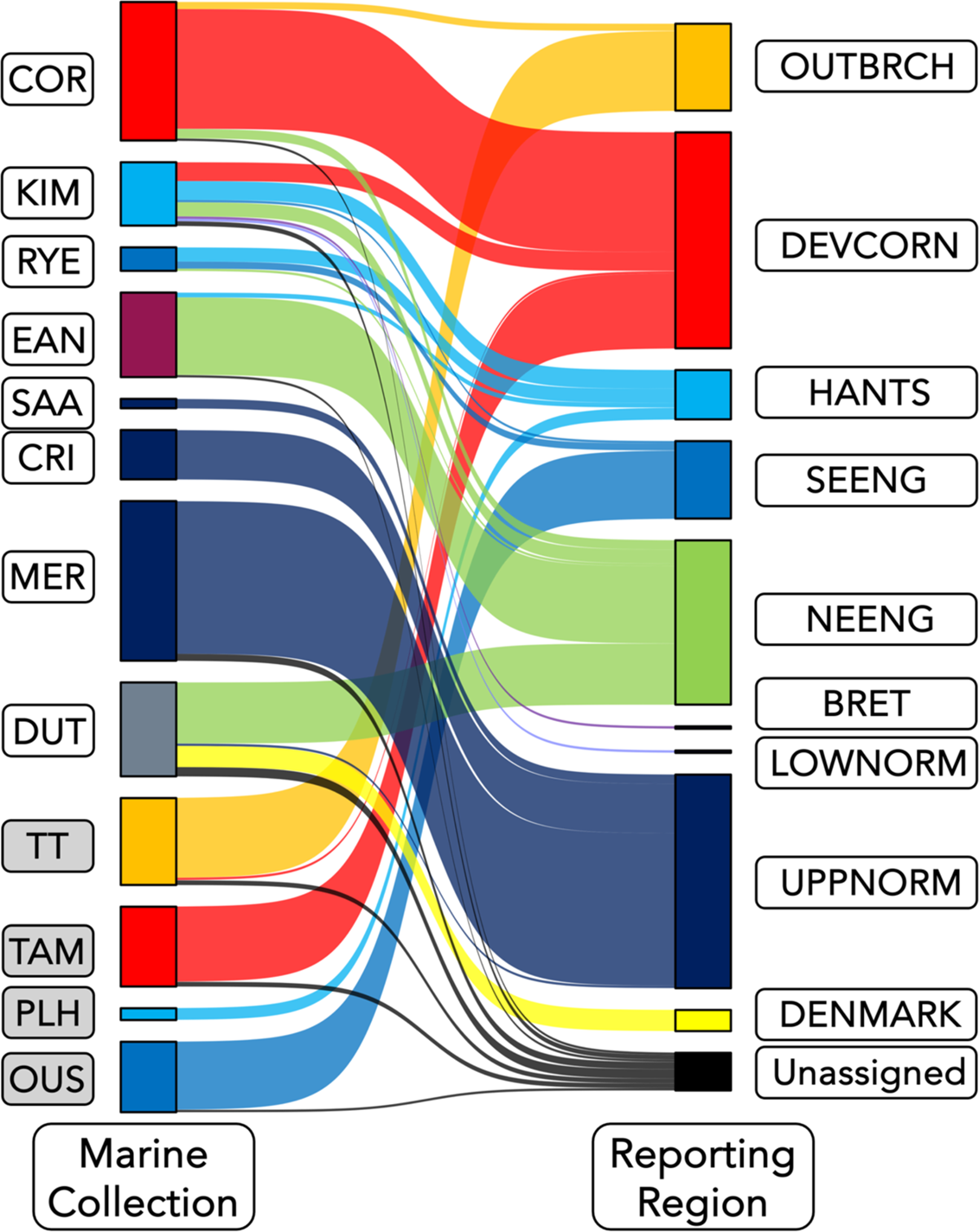
Sankey plot showing Individual Assignment of marine and estuarine caught anadromous trout to reporting region of origin. Marine and estuarine collections are colour coded by the reporting region they are located in while reporting regions are colour coded as given in Figure 1. Individuals were considered ‘Unassigned’ if the maximum probability of assignment to any reporting group was < 0.7. Marine collection abbreviations: COR – southern Cornwall; KIM – Kimmeridge Bay; RYE – Rye Harbour; EAN – East Anglian drift-net fishery; SAA – Saâne illegal nets; CRI – Criel-sur-Mer recreational beach nets; MER – Mers-les-Bains and Le Tréport recreational beach nets; DUT – Dutch commercial fishery by-catch. Estuarine collection abbreviations: TT – Taw/Torridge shared estuary; TAM – River Tamar tidal limit fish trap; PLH – Poole Harbour; OUS – Sussex Ouse estuary recreational rod fishery. Reporting group abbreviations: OUTBRCH – outer Bristol Channel; DEVCORN – Devon and Cornwall; HANTS – Hampshire Basin; SEENG – southeast England; NEENG – northeast England; BRET – Bretagne; LOWNORM – Lower Normandy; UPPNORM – Upper Normandy; DENMARK – Denmark.

The marine collections were more variable in their assignments to reporting group (Figure 3 and 4). Similar to the estuarine collections, some of the marine collections showed minimal variation in assignment outside of their expected reporting groups. For instance, sea trout in the collections from SAA, CRI and MER, which were caught in French waters in nets set close to the shore at the mouths of the Saâne, Yères and Bresle rivers, respectively, caught only fish from the UPPNORM reporting group (Figures 3 and 4). Likewise, in southwest England the COR sea trout samples were dominated by fish from the DEVCORN reporting group, with minor contributions from both OUTBRCH and NEENG rivers.

By contrast, the sea trout caught at KIM and RYE in southern England were more variable in their origins. Adult fish from six regions were caught at KIM, originating mainly from the three southern English reporting groups (DEVCORN, HANTS and SEENG). However, fish from BRET, LOWNORM and NEENG were also sampled here (Figures 3 and 4), while sea trout originating from the HANTS, SEENG and NEENG regions were sampled at RYE.

The two collections from the southern North Sea (EAN and DUT) were dominated by fish originating from the NEENG reporting group, with a significant contribution of trout from Danish rivers to the DUT samples. There were only minor contributions from English Channel reporting groups to these collections, with two fish of HANTS origin caught in the EAN nets and a single UPPNORM sea trout caught in Dutch waters (Figure 4).

### 3.5 Migration distances

Migration distances between the twelve marine and estuarine collections and the rivers of each reporting group are presented in Supplementary Table 6. This shows that the majority of sea trout were on average captured in close proximity to their natal rivers. For instance, the average capture distance for HANTS fish caught at KIM was 63.6 km. However, there are instances of very long-distance movements of sea trout, especially for those originating in NEENG rivers. THE NEENG fish caught at KIM and COR were on average 800 km and 965 km from their natal rivers (Supplementary Table 6).

## 4 Discussion

Here we present an extensive SNP-based genetic baseline for trout from English Channel and surrounding rivers, describing extensive, regional-based genetic structuring that allows high-confidence assignment of marine-caught sea trout to their region of origin.

### 4.1 Trout populations show strong regional genetic structure

The strong regional structuring of the trout populations in rivers screened here reiterates a pattern of distinct genetic groupings spanning sometimes long stretches of coastline and commonly observed in many anadromous salmonid species (Beacham et al., 2020; Beacham et al., 2021; Bradbury et al., 2015; Koljonen et al., 2014; Layton et al., 2020; Small et al., 2015). At the broadest scale, populations were split into two distinct eastern and western groups, with the split corresponding approximately with the Isle of Portland on the English side of the Channel and the Cotentin Peninsula on the French side. The Cotentin Peninsula and the relatively shallow waters to the north of the peninsula have previously been identified as a significant feature in the genetic structuring of a variety of marine organisms (Dauvin, 2012), including northern French trout populations (Quéméré et al., 2016).

Within each of the two main trout population groupings finer-scales of genetic structuring were also found. Three genetic groups of trout were identified in rivers entering the Channel on both the English and French Channel coasts. These corresponded with the three main geological zones existing on both sides of the Channel and it is likely that the genetic patterns observed are associated with the geology/water chemistry of the waters in which these fish live. Multiple, interacting factors help determine the chemical composition of river water. Of particular importance is underlying geology, which has a strong influence on pH, conductivity and concentrations of dissolved ions (Jarvie, Oguchi, & Neal, 2002; Liu, Weller, Correll, & Jordan, 2000; Rothwell et al., 2010). Brittany and southern Devon/Cornwall are dominated by Devonian age bedrock with granitic inclusions (e.g. the tors of Dartmoor), resulting in more acidic river water (pH≤7) with low conductivity.

Additionally, the upland areas of Brittany, Devon and Cornwall are dominated by blanket peat bog, reinforcing the acidic nature of river water in the area. Further east along both coasts in Normandy and south and southeast England the geology is dominated by Cretaceous era limestones and chalks, resulting in river water with pH values consistently above 7.

It has been suggested that the geological characteristics, and therefore chemical characteristics, of river catchments may be an important factor in determining the accuracy of homing through olfactory-based imprinting during smolting (Keefer & Caudill 2014), which may help to maintain regional structuring via reduced straying between genetically distinct groups of rivers (Bourret, Dionne, Kent, Lien, & Bernatchez, 2013). Additionally, underlying geology has been proposed to be a selective agent in the process of local adaptation in Atlantic salmon (Bourret et al., 2013). The hierarchical genetic structure detected here in English Channel trout also occurs in Atlantic salmon populations inhabiting rivers flowing into the Channel, with these patterns also having been linked to underlying geology (Ikediashi et al., 2018; Perrier, Guyomard, Bagliniere, & Evanno, 2011). Moreover, the locations of transitions in genetic profiles between groups are coincident in both species, providing stronger evidence that underlying geology is playing a major role in driving local adaptation in trout living along these coasts.

### 4.2 Consequences of regional structure for assignment to the baseline

The greater success of assignments to regions of origin reflects the metapopulation structure found in many salmonid species that have anadromous life-history stages (Schtickzelle & Quinn, 2007), with rivers in close proximity connected by gene flow via straying individuals from neighbouring rivers. Straying appears to be an integral part of salmonid life history. For instance, in a Danish fjord system, Källo *et al*. (2022) found high levels of straying of anadromous trout across multiple life history stages. Brown trout populations show strong regional genetic structuring (Bekkevold et al., 2020; Koljonen et al., 2014; Prodöhl et al., 2017), especially for rivers in the Channel region (King et al., 2016; King et al., 2020; Quéméré et al., 2016); within regional groups, however, there tend to be low levels of differentiation between populations in neighbouring rivers. For reporting groups with the largest sea trout runs (OUTBRCH, DEVCORN, NEENG, LOWNORM and UPPNORM) mean pairwise F_ST_ values were ≤0.04, indicative of little genetic differentiation between rivers within regions. Conversely, mean pairwise F_ST_ values between reporting groups were generally >0.08, supporting the assertion that genetic assignment performs better when there are large genetic distances between baseline stocks (Araujo, Candy, Beacham, White, & Wallace, 2014). Other salmonid fishery stock composition studies utilising extensive genetic baselines have also found greater assignment success to regional groups of geographically proximate rivers rather than to individual rivers (Bekkevold et al., 2021; Griffiths et al., 2010; Harvey et al., 2019; King et al., 2016; Koljonen et al., 2014; Prodöhl et al., 2017). In some cases, reporting groups have incorporated rivers covering from several hundreds to thousands of kilometres of coastline (Gilbey et al., 2016; Gilbey et al., 2018; Jeffery et al., 2018; Wennevik et al., 2019).

To minimise biases in estimates of stock composition, a reasonably complete baseline is necessary to capture the genetic signal of the potentially important stocks that may be present in mixtures (Araujo et al., 2014). One advantage for assignment studies is that the metapopulation structure often found in salmonid species (Schtickzelle & Quinn, 2007) reduces the need to sample all rivers potentially contributing to marine catches. It is not always possible, either logistically or financially, to exhaustively sample all sea trout-producing rivers in a region. Thus, a valid assumption of a regionally based assignment strategy is that samples originating from rivers not included in the baseline will likely be allocated to rivers from the same region, an approach that can reduce overall project costs (Beacham et al., 2020), albeit at the expense of a possible loss of finer resolution.

### 4.3 Stock structure of marine and estuarine collections

In the current study assignment results showed only very limited evidence of stock mixing of sea trout in the four estuarine collections. We can assume that these collections are the result of sampling local fish returning to their natal river prior to spawning. This was confirmed by the IA to river analyses (Supplementary Tables 5.09-5.12), which showed that the majority of fish caught in estuaries assigned to rivers flowing into the four estuaries. However, there were some fish that were clearly straying into these estuaries, with, for example, a NEENG fish caught in the recreational rod fishery in the Sussex Ouse (OUS), and three DEVCORN group fish caught in the net fishery in the Taw/Torridge (TT) estuary.

Similarly, four of the marine-caught collections (COR, SAA, CRI, MER) were predominantly sampling fish from local rivers. The main COR sampling sites were in Cawsand Bay, situated at the seaward edge of the Tamar estuary, with four major sea trout rivers (LYN, TAM, TAV, PLY) flowing out through the estuary. While few of the fish could reliably be assigned to river of origin, the main river-level assignments covered an ∼80 km stretch of coast within the DEVCORN reporting group from the East and West Looe rivers (25 km to the west of the estuary) to the Dart (∼55 km to the east of the estuary). Previous research has shown a degree of straying of sea trout from rivers along this stretch of coast into three of the Tamar estuary rivers (King et al., 2016). Likewise, the three samples of sea trout from the Upper Normandy coast (SAA, CRI, MER) also sampled predominately local fish. The nets in all three locations were recreational nets set from beaches during May to July when, again, fish would be returning to freshwater prior to spawning. Such targeting of local populations is not an uncommon feature of coastal fisheries targeting salmonids species. Fisheries for Atlantic salmon and Arctic charr on the Labrador coast of Canada (Bradbury et al., 2015; Bradbury et al., 2018; Layton et al., 2020) typically sampled fish from within ∼150 km of the capture site. Similarly, net fisheries for sea trout in the Gulf of Finland have been shown to be catching fish predominantly from rivers proximal to the netting areas (Koljonen et al., 2014).

### 4.4 Southern North Sea collections are dominated by NE English sea trout

The two marine collections from the southern North Sea (EAN and DUT) were dominated by fish from rivers in northeast England, i.e. the NEENG reporting group. The sea trout originating from rivers in this region are known to make long marine migrations, predominately migrating south along the east English North Sea coast. For instance, many sea trout tagged in the River Tweed have been caught in drift net fisheries along the East Anglian coast as well as in Dutch, German and Danish waters (Malcolm et al., 2010). This migration pattern has been confirmed using genetic assignment tests (Bekkevold et al., 2021). Thus, the southern North Sea appears to be important feeding grounds for multiple North Sea trout stocks (Bekkevold et al., 2021), with the results presented here providing evidence of sea trout originating from English Channel rivers (both English and French) also utilising this area.

### 4.5 Eastwards movements of southern English sea trout

The results for the KIM, RYE, EAN and DUT collections highlight a tendency for some of the sea trout from Channel rivers to move in an easterly direction once entering the marine environment. DEVCORN origin-fish were caught in Dorset at KIM and HANTS origin sea trout were present in the EAN collections and formed the majority of the fish sampled from RYE. Additionally, an UPPNORM fish was caught in the DUT net fishery. Previous historical tagging studies on sea trout smolts and kelts from the River Axe (DEVCORN reporting group) have shown that although on entering the marine environment the majority migrated west, some of the tag returns were from Hampshire rivers (HANTS) to the east, coastal nets along the Dorset and Hampshire coasts and the southern North Sea (Potter et al., 2017; Solomon, 1994). These fish appeared to be following the dominant west to east current that flows along the northern (English) side of the Channel into the southern North Sea (Dauvin, 2019; Winther & Johannessen, 2006).

### 4.6 Long-distance and cross-Channel movements

Some instances of very long-distance movements of sea trout from rivers in the NEENG reporting group were observed, with sea trout from northeast England being sampled from COR (4 fish), KIM (6 fish), RYE (1 fish). Additionally, a single sea trout caught in the Sussex Ouse recreational rod fishery had a probability (p=0.68) just below our 0.7 cut-off of originating from a river in the NEENG reporting group (Supplementary Table 5.12). Historic tagging studies undertaken on multiple life history stages of River Tweed sea trout have recorded only a single tag recovery from the English Channel (Malcolm et al., 2010). For the NEENG origin fish caught at Cawsand Bay, this represents a migration distance of ∼1000 km (Supplementary Table 6).

There were only two confirmed instances of cross-Channel movements of sea trout with individuals sampled at KIM originating from the BRET and LOWNORM reporting groups. Such cross-Channel movements do appear to be uncommon with only three tag recoveries from the northern French coast of sea trout tagged in southern English rivers (Potter et al., 2017). This finding is in contrast with the situation in the Irish Sea where frequent movements of trout from eastern Irish rivers into British coastal waters and *vice versa* were reported (Prodöhl et al., 2017).

### 4.7 Bycatch threats to sea trout during marine sojourns

In the marine environment sea trout exhibit a mainly piscivorous diet, with species such as sprat (*Sprattus sprattus*), sand eels (*Ammodytes* spp.) and herring (*Clupea harengus*) being dominant components of the diet (Knutsen, Knutsen, Gjosaeter, & Jonsson, 2001; Poiesz, Witte, & van der Veer, 2020; Roche et al., 2017). There are extensive commercial fisheries for two of these species (sprat and herring) in the southern North Sea and English Channel (Dauvin, 2019; Knijn, Boon, Heessen, & Hislop, 1993) and it is likely that there is widespread bycatch of sea trout in these fisheries, although bycatch levels appear to be under-recorded (Elliott et al., 2023). Additionally, it is likely that there will be bycatch in fisheries for fish species that have overlapping prey spectra with sea trout. For instance, our samples from RYE were caught in a net fishery that targets sea bass (*Dicentrarchus labrax*), which, like sea trout, are known to also feed on sprat and sand eel (Kelley, 1987; Spitz, Chouvelon, Cardinaud, Kostecki, & Lorance, 2013).

### 4.8 Management implications

The results presented here have implication for the management of sea trout in inshore waters around the English Channel and southern North Sea. Currently, for the UK, there is an extensive body of national and regional legislation designed to protect migratory salmonids from exploitation in inshore fisheries (Sumner, 2015); measures include protection from incidental capture in non-target fisheries and total netting bans in estuarine areas. However, some of these measures lack consistency across different regions. For instance, net headline – the recommended depth below which nets should be set – varies between 1.5 m and 3 m in different Inshore Fisheries & Conservation Authority regions along the southern English coast (Sumner, 2015).

Marine protected areas (MPAs) offer one route to safeguard sea trout during their marine migrations. Such areas offer protection within the designated region to both resident fish species and also species that transit through them (Breen, Posen, & Righton, 2015). At present, however, evidence that MPAs are effective for the conservation of highly mobile species such as sea trout is limited (Breen et al., 2015). Nevertheless, to determine the efficacy of MPAs, to regulate fisheries and contribute to policy we require knowledge of where and when individuals are at sea (O’Sullivan et al., 2022). Genetic assignment studies, such as that presented here can help identify both fish movements and fisheries pressure on species, thereby providing evidence crucial to the designation and meaningful placement of MPAs (Jeffery et al., 2022).

Effective conservation of sea trout stocks in the marine environment therefore must include measures to minimise the risk of incidental capture. Based on inter-river connectivity, as determined from population genetic data and prioritisation analyses, a number of potential MPAs for English Channel sea trout have recently been proposed, (Vanhove et al., submitted). Scenarios took into account factors such as fishing density and other human effects on the marine environment, resulting in proposed protection areas along the south Devon and Cornish coasts, northern Brittany, Lower Normandy, the area between Dorset/Hampshire and the Cotentin Peninsula and the eastern Channel between Kent/Sussex and Upper Normandy (Vanhove et al., submitted). Interestingly, two of these areas (Dorset/Hampshire and Kent/Sussex) are where we found the highest levels of stock mixing in our marine sea trout samples, strengthening the evidence that these areas should be designated as protected areas for sea trout in the English Channel.

## Acknowledgements

We thank Simon Toms, Lawrence Talks, Willie Roche and the staff of the Environment Agency, Devon & Severn, Southern and Cornwall Inshore Fisheries Conservation Authorities, Inland Fisheries Ireland, INRAE U3E Unit, Office Français de la Biodiversité, Bretagne Grands Migrateurs, Seinormigr, FDPPMA14, 22, 27, 29, 35, 50, 62, 76 & 80 and André Breukelaar (Rijkswaterstaat) for co-ordinating and supporting sample collection.

## Funding Information

This research was funded by the European Union Interreg France (Channel) England programme project ‘Salmonid Management Around the Channel’ (SAMARCH) with additional funding from the Missing Salmon Alliance. DRO was supported by a GW4 NERC Doctoral Training Programme PhD studentship as part of the FRESH programme.

## Notes

### Competing Interest Statement

The authors have declared no competing interest.

