## Supplementary Materials for "Leveraging the genetic diversity of trout in the rivers of the British Isles and northern France to understand the movements of sea trout (*Salmo trutta* L.) around the English Channel"

**Index**

**Page 2** – SNP Baseline testing – methods

**Page 3** – SNP Baseline testing – results

**Page 5** – Details of marine and estuarine collections

**Page 8** – Supplementary Table 1 – Details of sampled baseline rivers

**Page 13** – Supplementary Table 6 – Least cost distances between marine and estuarine sampling locations and rivers in each reporting group

**Page 15 –** Supplementary Figure 2 – map of marine and estuarine sampling locations

**Page 16** – Supplementary Figure 2 – *RUBIAS* baseline group-level Leave-One-Out accuracy and efficiency

**Page 17** – Supplementary Figure 3 – *RUBIAS* baseline river-level Leave-One-Out accuracy and efficiency

**Page 18** – Supplementary Figure 4 – Comparison of mixed stock and individual assignment analyses for cBayes and *RUBIAS* for known-origin trout

**Page 19** – Supplementary Figure 5 – Evanno et al. (2005) delta *K* (Δ*K* ) results for the hierarchical STRUCTURE analyses of genetic structuring

**Page 20** – Supplementary references

**SNP Baseline testing - methods**

We used two widely utilised assignment software for mixed stock analyses (MSA) and individual assignment (IA) to both individual river and each of 14 reporting groups. *RUBIAS* uses a Bayesian conditional genetic stock identification model to provide mixture proportion estimates and assign individuals to population/stock of origin (Moran & Anderson 2019). Assignment proportions and their 95% credible intervals were generated using the Markov chain Monte Carlo (MCMC) method based on 100000 sweeps following a burn-in of 10000 sweeps. cBayes (Neaves *et al*. 2005) implements the Bayesian procedures of Pella and Masuda (2001). For stock composition estimation, eight 50 000-interation Markov Chain Monte Carlo (MCMC) chains were run, with initial values set at 0.9 for each chain for different samples. Means and 95% confidence intervals of the estimated stock contributions were determined from the combined final 1000 iterations from each chain. cBayes only performs IA to individual rivers. For IA to group we used an ‘allocate and sum’ approach (Wood *et al*. 1987).

Initial baseline testing was undertaken using a Leave-One-Out (LOO) analysis, performed using *RUBIAS*, to assess assignment accuracy (number of correctly assigned individuals divided by the total number assigned to each assignment unit) and efficiency (number of correctly assigned individuals divided by the total number known *a priori* for that assignment unit) at both reporting group (14 groups as defined by a neighbour-joining dendrogram - see main Results section) and river levels. To minimise potential type I errors, an assignment was assumed to be correct if the individual assignment probability was ≥ 0.7 (Vähä *et al*. 2011, Prodöhl et al. 2017, Bradbury *et al*. 2018).

A more realistic assessment of baseline performance is to use samples of known origin. Mixed stock and individual assignment analyses of 435 known origin individuals from 25 baseline rivers (Supplementary Table 1) was undertaken using both cBayes and *RUBIAS*, as described above. We calculated the proportion of correct assignment as the number of assignments with p ≥ 0.7 divided by the total number of individuals in each collection.

**SNP Baseline testing - results**

The Leave-One-Out analyses showed generally high assignment success to reporting group, with the majority of groups having >85% assignment accuracy and efficiency (Supplementary Figure 2). Conversely, assignment accuracy and efficiency to individual rivers was highly variable (Supplementary Figure 3). For some rivers, such as the Tyne (TYN), Severn (SEV) and Tresillian (TRE), both assignment accuracy and efficiency were in excess of 90% (Supplementary Figure 3), however for the majority of rivers, both accuracy and efficiency were much lower. For instance, for many of the rivers in the DEVCORN reporting group, accuracy and efficiency of assignment were well below 50% (Supplementary Figure 3).

Supplementary Figure 4 summarises the MSA and IA of known origin collections to river and reporting group of origin. Full results are presented in Supplementary Tables 2 and 3. Similar to the results for the LOO analysis, the majority of collections and individuals assigned to their known group of origin (Supplementary Figure 4) with high assignment probability/success. The exception was the collection from the Rother (ROT), with three individuals assigning with variable probabilities to the Lower Normandie group and River Orne (Supplementary Figure 4, Supplementary Tables 2 and 3). Conversely, assignment success to river of origin was variable between collections, with some showing high self-assignment *i.e.* ECL, TOR & SEU (Supplementary Figure 4). However, in general, assignment success was poor with two rivers (FOW & TEI) showing zero self-assignment success.

It should also be noted that there were differences in the ability of cBayes and *RUBIAS* to assign known-origin fish and collections to their river and reporting group of origin (Supplementary Figure 4). In general, across the 25 collections, cBayes was marginally more successful than *RUBIAS* at assigning individuals and collections to river and reporting group of origin (Supplementary Table 2). There were, however, clear differences in assignment success between the two software. For the TAM, cBayes assigned all individuals to the Tamar, while only eight (out of 19) individuals were assigned to the Tamar by RUBIAS. Conversely, *RUBIAS* was better at self-assigning individuals from the Nevern (NEV) than cBayes (Supplementary Figure 4).

**Details of marine and estuarine collections**

**COR - South Cornwall**

The majority of sea trout sampled from south Cornwall were caught by a commercial netsman fishing in Whitsand Bay under license exemptions for the SAMARCH project in 2018, 2019 & 2021. Additional samples were collected in 2010-2013 as part of a previous project by recreational anglers targeting sea bass from Queener Point at the eastern end of Whitsand Bay. An additional fish was caught by a recreational angler targeting sea bass at Porthallow, Cornwall. Final sample size was 59 individuals.

**KIM - Kimmeridge**

All sea trout sampled from Kimmeridge Bay were caught by a commercial netsman fishing under license exemptions for the SAMARCH project in 2018, 2019 & 2021. Final sample size was 27 individuals.

For both South Cornwall and Kimmeridge Bay, fixed surface-set 200m multi monofilament gill nets were used. Each net was anchored to the seabed by heavy weights with marker buoys on either end with led lights as a warning. Mesh sizes used ranged between 9.50 cm and 11.40 cm and nets were between 4.08 and 5.70 m in height when set. Nets were set in the evening, near to (between 20m to 1000m), and mainly perpendicular to the shore, stretched between two fixed points and were recovered the following morning. On each sampling trip three or four nets were deployed.

**RYE - Rye Harbour**

Sea trout were caught in 2020 and 2021 by a commercial fisherman operating nets targeting a mixed assemblage of species. An additional sample was caught by a recreational fisherman off Seaford Head, East Sussex in 2010. Final sample size was 10 individuals.

**EAN - East Anglia**

The East Anglian net fishery is a commercial fishery specifically targeting migratory salmonids. Samples were obtained from multiple locations along the Norfolk coast. The precise capture location of some fish is not known. The fishery utilises a mix of drift and fixed nets. The fixed nets are gillnets that are usually pinned out in sandy bays at low tide. They catch through the flooding tide and any catch is collected when the tide drops again. Final sample size was 36 individuals.

**DUT - Netherlands**

Sea trout samples were collected from commercial fishers operating along the Dutch North- and Wadden Sea coasts between Hoek van Holland and Lauwersoog. These fishers were contracted to provide salmonid fish samples for a range of research projects funded by Rijkswaterstaat (RWS), the Implementing Body of the Ministry of Transport, Public Works and Water Management, in the Netherlands. The nets used were a combination of fyke- and bagnets. Final sample size was 40 individuals.

**SAA & MER - North France**

Four scale sets were available from sea trout caught in June 2021 in illegally-set fixed nets set just outside the estuary of the Saâne river.

Scale samples from Criel-sur-Mer at the mouth of the Yères and Mers-les-Bains & Le Tréport just north and south, respectively of the mouth of the Bresle, were collected as part of a survey of legal recreational net fishing between 1984 and 1994. These consists of 50m nets that were set perpendicular to the coast, usually not far from estuaries (Fagard & Beaulaton 2018). Because of their geographical proximity, the samples for Mers-les-Bains and Le Tréport were combined. Final sample size was 21 and 68 individuals for Criel-sur-Mer and Mers-les-Bains/Le Tréport, respectively.

**TT - Taw/Torridge estuary (Appledore)**

The majority of the sea trout were sampled from commercial estuarine nets specifically targeting migratory salmonids. The netting locations were at Appledore (51.057167, -4.193025) at the confluence of the Taw and Torridge rivers. The scale samples were collected in 2011 as part of a previous project and reanalysed for this project. Additionally, three scale sets were obtained from sea trout caught at coastal locations in close proximity to the Taw/Torridge estuary during netting activities undertaken specifically for the SAMARCH project by a commercial netsman. Final sample size was three individuals from coastal locations and 34 from the Appledore nets.

**TAM – Gunnislake Fish Trap**

Adult sea trout were caught during their upstream spawning migration in a purpose-designed fish trap associated with a fish pass located on a weir at the tidal limit at Gunnislake, Cornwall (grid reference 50.519, −4.206). The River Tamar is one of three Environment Agency Index catchments and is subjected to intensive monitoring to both inform and improve wider management of migratory salmonids in England, including extensive juvenile electrofishing surveys, tagging of smolts during spring migration, and trapping of returning adults. The final sample size was 34, taken from fish collected in 2017 & 2020.

**PLH – Poole Harbour**

Scale samples were collected from sea bass nets set within Poole Harbour. The final sample size was five individuals.

**OUS - Sussex Ouse Recreational Fishery**

The Sussex Ouse sea trout analysed in this study were caught between 2010 and 2015 by recreational anglers from the Ouse Angling Preservation Society in waters downstream of the tidal limit of the river at Barcombe Mills (50.915317, 0.034699). A small number were captured further downstream at Lewes (Cowlease Farm – 50.902246, 0.020193). The open season for sea trout fishing on R. Ouse is from 1 May until 31 October. Final sample size was 30 individuals.

**Supplementary Table 1** – Details of rivers sampled for resident brown trout for the construction of the single nucleotide polymorphism baseline. Latitude and longitude data are given for the mouth of each river.

| **River** | **Code** | **Country** | **Reporting Group** | **Latitude** | **Longitude** | **Number genotyped** | **Removed Full sibs** | **Failed samples** | **Baseline samples** | **Known origin samples** |
| --- | --- | --- | --- | --- | --- | --- | --- | --- | --- | --- |
| Arigideen | ARI | Ireland | South east Ireland | 51.625845 | -8.668346 | 20 | 0 | 0 | 20 | 0 |
| Colligan | COL | Ireland | South east Ireland | 52.071162 | -7.513121 | 20 | 0 | 0 | 20 | 0 |
| Sow | SOW | Ireland | South east Ireland | 52.316017 | -6.363038 | 20 | 2 | 0 | 18 | 0 |
| Owenavaragh | QWE | Ireland | South east Ireland | 52.654237 | -6.220131 | 20 | 0 | 0 | 20 | 0 |
| Inch | INC | Ireland | South east Ireland | 52.711022 | -6.166956 | 20 | 0 | 0 | 20 | 0 |
| Avoca | AVO | Ireland | South east Ireland | 52.793996 | -6.135749 | 41 | 0 | 0 | 20 | 21 |
| Nevern | NEV | Wales | Outer Bristol Channel | 52.026942 | -4.85074 | 40 | 0 | 1 | 30 | 9 |
| West Cleddau | WCL | Wales | Outer Bristol Channel | 51.663269 | -5.154108 | 32 | 0 | 0 | 32 | 0 |
| East Cleddau | ECL | Wales | Outer Bristol Channel | 51.663269 | -5.154108 | 40 | 0 | 0 | 30 | 10 |
| Taff | TAF | Wales | Outer Bristol Channel | 51.7168 | -4.41441 | 30 | 5 | 0 | 25 | 0 |
| Twyi | TWY | Wales | Outer Bristol Channel | 51.7168 | -4.41441 | 32 | 0 | 1 | 31 | 0 |
| Tawe | TWE | Wales | Outer Bristol Channel | 51.610081 | -3.927836 | 30 | 5 | 1 | 24 | 0 |
| Ogmore | OGM | Wales | Outer Bristol Channel | 51.468126 | -3.645016 | 24 | 0 | 0 | 24 | 0 |
| Wye | WYE | Wales | Inner Bristol Channel | 51.608194 | -2.662412 | 30 | 0 | 0 | 30 | 0 |
| Severn | SEV | England | Inner Bristol Channel | 51.657636 | -2.589118 | 29 | 0 | 1 | 28 | 0 |
| Bristol Avon | BAV | England | Inner Bristol Channel | 51.506059 | -2.726962 | 30 | 1 | 0 | 29 | 0 |
| Doniford | DON | England | Outer Bristol Channel | 51.184262 | -3.304221 | 24 | 1 | 0 | 23 | 0 |
| East Lyn | ELY | England | Outer Bristol Channel | 51.23501 | -3.829387 | 30 | 3 | 0 | 27 | 0 |
| Taw | TAW | England | Outer Bristol Channel | 51.07585 | -4.239722 | 30 | 0 | 0 | 30 | 0 |
| Torridge | TOR | England | Outer Bristol Channel | 51.07585 | -4.239722 | 38 | 0 | 1 | 29 | 8 |
| Camel | CAM | England | Devon & Cornwall | 50.577934 | -4.948684 | 64 | 0 | 0 | 40 | 24 |
| Menalhyl | MEN | England | Devon & Cornwall | 50.468438 | -5.039764 | 24 | 0 | 0 | 24 | 0 |
| Red River | RED | England | Land's End | 50.230971 | -5.392895 | 30 | 0 | 3 | 27 | 0 |
| Hayle | HAY | England | Land's End | 50.200946 | -5.437558 | 32 | 2 | 0 | 30 | 0 |
| Trevaylor | TRV | England | Land's End | 50.125869 | -5.524697 | 25 | 0 | 0 | 25 | 0 |
| Crowlas | CRO | England | Land's End | 50.125802 | -5.480418 | 25 | 0 | 0 | 25 | 0 |
| Tresillian | TRE | England | Devon & Cornwall | 50.142221 | -5.029527 | 30 | 0 | 0 | 30 | 0 |
| Fal | FAL | England | Devon & Cornwall | 50.142221 | -5.029527 | 30 | 0 | 0 | 30 | 0 |
| Fowey | FOW | England | Devon & Cornwall | 50.323694 | -4.642354 | 50 | 0 | 0 | 32 | 18 |
| West Looe | WLO | England | Devon & Cornwall | 50.350145 | -4.44909 | 30 | 0 | 0 | 30 | 0 |
| East Looe | ELO | England | Devon & Cornwall | 50.350145 | -4.44909 | 40 | 0 | 1 | 29 | 10 |
| Seaton | SEA | England | Devon & Cornwall | 50.362856 | -4.388246 | 33 | 0 | 1 | 32 | 0 |
| Lynher | LYN | England | Devon & Cornwall | 50.32216 | -4.152776 | 30 | 0 | 1 | 29 | 0 |
| Tamar | TAM | England | Devon & Cornwall | 50.32216 | -4.152776 | 56 | 0 | 1 | 36 | 19 |
| Tavy | TAV | England | Devon & Cornwall | 50.32216 | -4.152776 | 30 | 0 | 1 | 29 | 0 |
| Plym | PLY | England | Devon & Cornwall | 50.32216 | -4.152776 | 30 | 0 | 1 | 29 | 0 |
| Yealm | YEA | England | Devon & Cornwall | 50.300346 | -3.954329 | 30 | 0 | 0 | 30 | 0 |
| Erme | ERM | England | Devon & Cornwall | 50.310927 | -4.073074 | 59 | 0 | 0 | 37 | 22 |
| Devon Avon | DAV | England | Devon & Cornwall | 50.276325 | -3.88919 | 51 | 1 | 1 | 28 | 21 |
| Dart | DAR | England | Devon & Cornwall | 50.333009 | -3.555909 | 72 | 0 | 0 | 42 | 30 |
| Teign | TEI | England | Devon & Cornwall | 50.539001 | -3.492309 | 38 | 0 | 0 | 30 | 8 |
| Exe | EXE | England | Devon & Cornwall | 50.608556 | -3.41592 | 61 | 0 | 0 | 32 | 29 |
| Otter | OTT | England | Devon & Cornwall | 50.627911 | -3.303624 | 30 | 0 | 0 | 30 | 0 |
| Axe | AXE | England | Devon & Cornwall | 50.701426 | -3.055595 | 30 | 0 | 1 | 29 | 0 |
| Brit | BRI | England | Hampshire Basin | 50.707775 | -2.764488 | 30 | 0 | 1 | 29 | 0 |
| Frome | FRO | England | Hampshire Basin | 50.677329 | -1.936547 | 78 | 0 | 0 | 48 | 30 |
| Piddle | PID | England | Hampshire Basin | 50.677329 | -1.936547 | 30 | 0 | 0 | 30 | 0 |
| Dorset Stour | DST | England | Hampshire Basin | 50.721962 | -1.735832 | 30 | 0 | 0 | 30 | 0 |
| Hampshire Avon | HAV | England | Hampshire Basin | 50.721962 | -1.735832 | 46 | 0 | 0 | 30 | 16 |
| Lymington | LYM | England | Hampshire Basin | 50.741284 | -1.512443 | 42 | 0 | 0 | 30 | 12 |
| Beaulieu | BEA | England | Hampshire Basin | 50.78117 | -1.374305 | 30 | 1 | 2 | 27 | 0 |
| Itchen | ITC | England | Hampshire Basin | 50.86682 | -1.366846 | 30 | 0 | 0 | 30 | 0 |
| Test | TES | England | Hampshire Basin | 50.86682 | -1.366846 | 42 | 5 | 0 | 25 | 12 |
| Hamble | HAM | England | Hampshire Basin | 50.843933 | -1.314478 | 30 | 5 | 0 | 25 | 0 |
| Meon | MEO | England | Hampshire Basin | 50.816412 | -1.241401 | 30 | 1 | 0 | 29 | 0 |
| Wallington | WAL | England | Hampshire Basin | 50.775205 | -1.108099 | 30 | 0 | 0 | 30 | 0 |
| Eastern Yar | YAR | England | South east England | 50.69769 | -1.094026 | 30 | 0 | 0 | 30 | 0 |
| Arun | ARU | England | South east England | 50.798915 | -0.540928 | 29 | 0 | 0 | 29 | 0 |
| Adur | ADU | England | South east England | 50.823797 | -0.247195 | 30 | 0 | 0 | 30 | 0 |
| Sussex Ouse | OUS | England | South east England | 50.776616 | 0.060959 | 42 | 0 | 2 | 31 | 9 |
| Eastern Rother | ROT | England | South east England | 50.925445 | 0.777134 | 40 | 0 | 0 | 30 | 10 |
| Dour | DOU | England | South east England | 51.111552 | 1.329468 | 30 | 0 | 0 | 30 | 0 |
| Great Stour | GST | England | South east England | 51.318806 | 1.383149 | 30 | 7 | 0 | 23 | 0 |
| Medway | MED | England | Thames & East Anglia | 51.462076 | 0.745395 | 30 | 0 | 1 | 29 | 0 |
| Thames | THA | England | Thames & East Anglia | 51.485119 | 0.795208 | 34 | 7 | 4 | 23 | 0 |
| Belstead Brook | BEL | England | Thames & East Anglia | 51.896767 | 1.358933 | 30 | 10 | 0 | 20 | 0 |
| Wensum | WEN | England | Thames & East Anglia | 52.572361 | 1.745058 | 25 | 1 | 1 | 23 | 0 |
| Glaven | GLA | England | Thames & East Anglia | 52.983507 | 0.96992 | 35 | 2 | 0 | 33 | 0 |
| Nar | NAR | England | Thames & East Anglia | 52.835396 | 0.341324 | 20 | 0 | 1 | 19 | 0 |
| Yorkshire Esk | ESK | England | North east England | 54.493882 | -0.611669 | 16 | 0 | 0 | 16 | 0 |
| Tyne | TYN | England | North east England | 55.010573 | -1.394313 | 52 | 0 | 0 | 32 | 20 |
| Coquet | COQ | England | North east England | 55.339596 | -1.567247 | 22 | 2 | 0 | 20 | 0 |
| Tweed | TWEE | England | North east England | 55.763122 | -1.981566 | 24 | 0 | 0 | 24 | 0 |
| Aber Wrac'h | ABE | France | Bretagne | 48.610399 | -4.582648 | 26 | 1 | 0 | 25 | 0 |
| Flèche | FLE | France | Bretagne | 48.644957 | -4.299911 | 31 | 2 | 0 | 29 | 0 |
| Kérallé | KER | France | Bretagne | 48.657244 | -4.228475 | 32 | 7 | 0 | 25 | 0 |
| Guillec | GUI | France | Bretagne | 48.688319 | -4.073144 | 31 | 4 | 1 | 26 | 0 |
| Horn | HOR | France | Bretagne | 48.688681 | -4.059893 | 32 | 0 | 2 | 30 | 0 |
| Yar | FYA | France | Bretagne | 48.674915 | -3.583281 | 32 | 3 | 1 | 28 | 0 |
| Léguer | LEG | France | Bretagne | 48.735102 | -3.560884 | 31 | 0 | 0 | 31 | 0 |
| Ic | IC | France | Bretagne | 48.600973 | -2.814563 | 31 | 1 | 0 | 30 | 0 |
| Frémur | FRE | France | Bretagne | 48.621228 | -2.143971 | 28 | 0 | 0 | 28 | 0 |
| Couesnon | COU | France | Bretagne | 48.637091 | -1.51632 | 35 | 3 | 0 | 32 | 0 |
| Selune | SEL | France | Bretagne | 48.648012 | -1.447503 | 33 | 8 | 1 | 24 | 0 |
| Sée | SEE | France | Bretagne | 48.648012 | -1.447503 | 35 | 4 | 0 | 25 | 6 |
| Sienne | SIE | France | Bretagne | 48.990324 | -1.569532 | 34 | 2 | 0 | 32 | 0 |
| Saire | SAI | France | Bretagne | 49.609818 | -1.250799 | 30 | 0 | 0 | 30 | 0 |
| Vire | VIR | France | Lower Normandie | 49.365329 | -1.129752 | 31 | 2 | 0 | 29 | 0 |
| Seulles | SEU | France | Lower Normandie | 49.337897 | -0.456177 | 40 | 3 | 1 | 27 | 9 |
| Orne | ORN | France | Lower Normandie | 49.293151 | -0.246018 | 33 | 5 | 1 | 27 | 0 |
| Dives | DIV | France | Lower Normandie | 49.306222 | -0.095141 | 36 | 3 | 0 | 33 | 0 |
| Touques | TOU | France | Lower Normandie | 49.371122 | 0.068543 | 33 | 0 | 12 | 21 | 0 |
| Seine | SEI | France | Lower Normandie | 49.431671 | 0.084352 | 60 | 4 | 2 | 30 | 24 |
| Valmont | VAL | France | Upper Normandie | 49.765316 | 0.362524 | 30 | 3 | 3 | 24 | 0 |
| Durdent | DUR | France | Upper Normandie | 49.856446 | 0.607286 | 26 | 0 | 0 | 26 | 0 |
| Saâne | SAA | France | Upper Normandie | 49.906918 | 0.929678 | 30 | 0 | 1 | 29 | 0 |
| Scie | SCI | France | Upper Normandie | 49.918092 | 1.031633 | 28 | 0 | 0 | 28 | 0 |
| Arques | ARQ | France | Upper Normandie | 49.936621 | 1.082478 | 76 | 1 | 2 | 40 | 33 |
| Yères | YER | France | Upper Normandie | 50.033002 | 1.310304 | 27 | 0 | 0 | 27 | 0 |
| Bresle | BRE | France | Upper Normandie | 50.066402 | 1.368578 | 67 | 0 | 5 | 36 | 26 |
| Somme | SOM | France | Upper Normandie | 50.23594 | 1.523448 | 31 | 1 | 3 | 27 | 0 |
| Authie | AUT | France | Upper Normandie | 50.374055 | 1.566111 | 14 | 0 | 0 | 14 | 0 |
| Canche | CAN | France | Upper Normandie | 50.548323 | 1.578664 | 27 | 2 | 6 | 19 | 0 |
| Liane | LIA | France | Upper Normandie | 50.737078 | 1.580767 | 34 | 1 | 0 | 33 | 0 |
| Slack | SLA | France | Upper Normandie | 50.805 | 1.600931 | 33 | 4 | 0 | 29 | 0 |
| Sneum | SNE | Denmark | Denmark | 55.905366 | 8.355854 | 40 | 0 | 2 | 38 | 0 |
| Skjern | SKJ | Denmark | Denmark | 55.361061 | 8.334486 | 29 | 0 | 0 | 29 | 0 |
| Hatchery J | HATJ | Hatchery | French hatchery | - | - | 30 | 0 | 0 | 30 | 0 |
| Hatchery PF | HATPF | Hatchery | French hatchery | - | - | 26 | 0 | 0 | 26 | 0 |
| Total |  |  |  |  |  | 3699 | 125 | 71 | 3067 | 436 |

**Supplementary Table 6** Least cost distances between marine and estuarine sampling locations and rivers in each reporting group. Distances were calculated using the marmap R package (Pante & Simon-Bouhet, 2013). Min – minimum distance between sea trout sampling location and a river in a reporting group; max - maximum distance between sea trout sampling location and a river in a reporting group; average - average distance between sea trout sampling location and each river in a reporting group. Distances are presented only for the reporting groups to which sea trout from each marine or estuarine collection were assigned, based on Individual Assignment results (Supplementary Tables 5.01-5.12).

|  |  | **Marine** | | | | | | | | **Estuarine** | | | |
| --- | --- | --- | --- | --- | --- | --- | --- | --- | --- | --- | --- | --- | --- |
| **Reporting Group** | **Distance** | **COR** | **KIM** | **RYE** | **EAN** | **SAA** | **CRI** | **MER** | **DUT** | **TT** | **TAM** | **PLH** | **OUS** |
| OUTBRCH | min | 288 | - | - | - | - | - | - | - | 0 | - | - | - |
|  | max | 363 | - | - | - | - | - | - | - | 92 | - | - | - |
|  | average | 330.3 | - | - | - | - | - | - | - | 61.3 | - | - | - |
| DEVCORN | min | 7 | 76 | - | - | - | - | - | - | 81 | 0 | - | - |
|  | max | 209 | 372 | - | - | - | - | - | - | 397 | 215 | - | - |
|  | average | 71.9 | 176.7 | - | - | - | - | - | - | 287.1 | 70.7 | 197.9 | - |
| HANTS | min | - | 30 | 144 | 418 | - | - | - | - | - | - | 0 | - |
|  | max | - | 86 | 272 | 546 | - | - | - | - | - | - | 77 | - |
|  | average | - | 63.6 | 181.3 | 455.1 | - | - | - | - | - | - | 44.6 | - |
| SEENG | min | - | 90 | 6 | - | - | - | - | - | - | - | - | 0 |
|  | max | - | 271 | 143 | - | - | - | - | - | - | - | - | 107 |
|  | average | - | 172.0 | 73.7 | - | - | - | - | - | - | - | - | 53.8 |
| NEENG | min | 879 | 715 | 491 | 225 | - | - | - | 515 | - | - | - | - |
|  | max | 1040 | 876 | 652 | 386 | - | - | - | 640 | - | - | - | - |
|  | average | 964.8 | 800.8 | 576.3 | 310.3 | - | - | - | 583.5 | - | - | - | - |
| BRET | min | - | 140 | - | - | - | - | - | - | - | - | - | - |
|  | max | - | 292 | - | - | - | - | - | - | - | - | - | - |
|  | average | - | 238.0 | - | - | - | - | - | - | - | - | - | - |
| LOWNORM | min | - | 161 | - | - | - | - | - | - | - | - | - | - |
|  | max | - | 215 | - | - | - | - | - | - | - | - | - | - |
|  | average | - | 199.2 | - | - | - | - | - | - | - | - | - | - |
| UPPNORM | min | - | - | - | - | 0 | 0 | 0 | 500 | - | - | - | - |
|  | max | - | - | - | - | 116 | 94 | 91 | 644 | - | - | - | - |
|  | average | - | - | - | - | 53.8 | 48.6 | 49.1 | 575.6 | - | - | - | - |
| DENMARK | min | - | - | - | - | - | - | - | 215 | - | - | - | - |
|  | max | - | - | - | - | - | - | - | 278 | - | - | - | - |
|  | average | - | - | - | - | - | - | - | 246.5 | - | - | - | - |

**Supplementary Figure 1** Map giving the approximate location for eight marine (white label) and four estuarine (grey label) collections of anadromous trout. Marine collection abbreviations: COR – southern Cornwall; KIM – Kimmeridge Bay; RYE – Rye Harbour; EAN – East Anglian drift-net fishery; SAA – Saâne illegal nets; CRI – Criel-sur-Mer recreational beach nets; MER – Mers-les-Bains and Le Tréport recreational beach nets; DUT – Dutch commercial fishery by-catch. Estuarine collection abbreviations: TT – Taw/Torridge shared estuary; TAM – River Tamar tidal limit fish trap; PLH – Poole Harbour; OUS – Sussex Ouse estuary recreational rod fishery

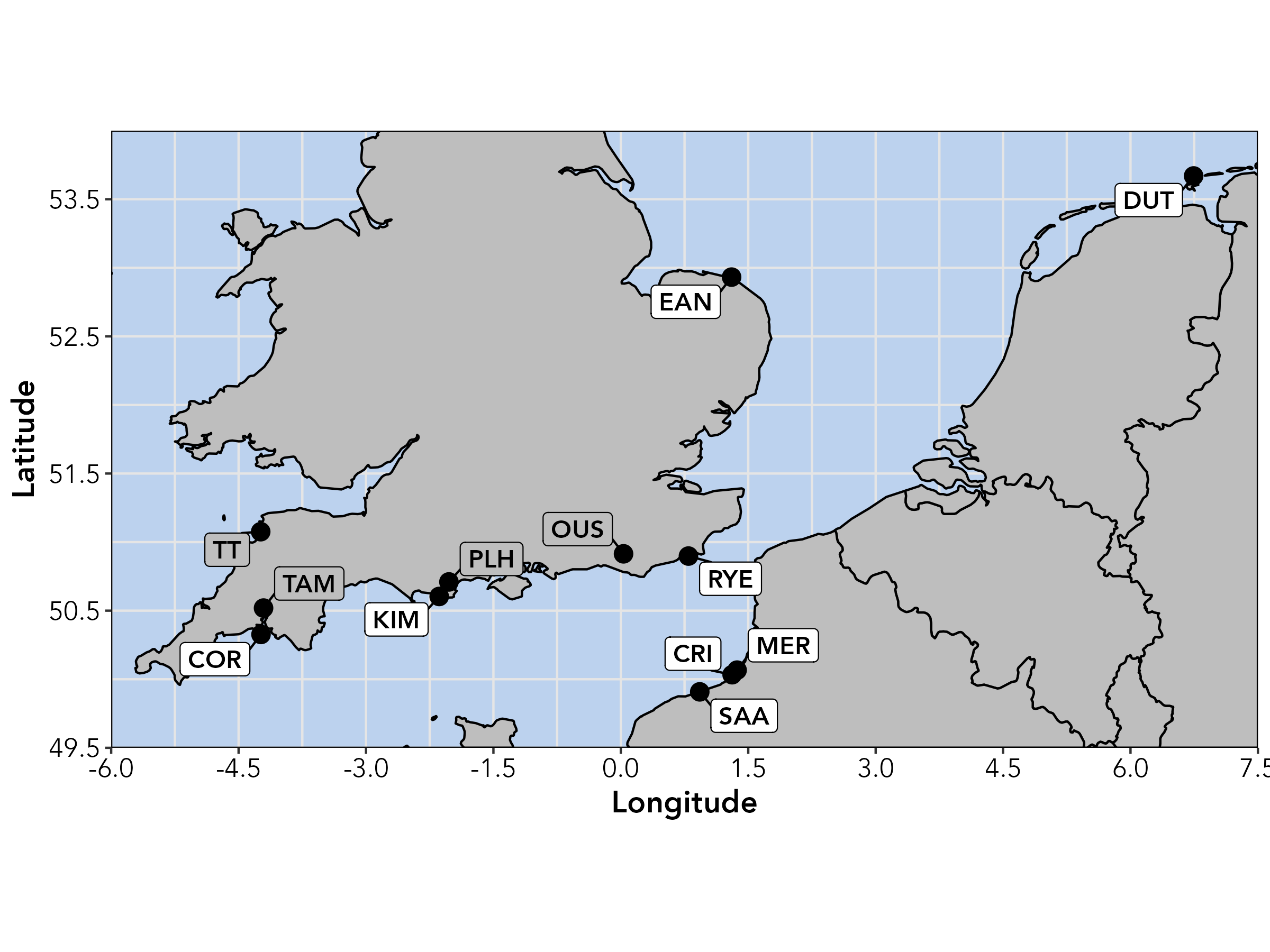

**Supplementary Figure 2** *RUBIAS* group-level Leave-One-Out accuracy (red) and efficiency (blue). Reporting group codes are SEIRE – south east Ireland, OUTBRCH – outer Bristol Channel, INNBRCH – inner Bristol Channel, DEVCORN – Devon & Cornwall, LANDSEND – Land’s End, HANTS – Hampshire Basin, SEENG – south east England, THAMESEA – Thames & East Anglia, NEENG – north east England, BRET – Bretagne, LOWNORM – Lower Normandie, UPPNORM – Upper Normandie, DENMARK – Denmark and FRHAT – French hatcheries.

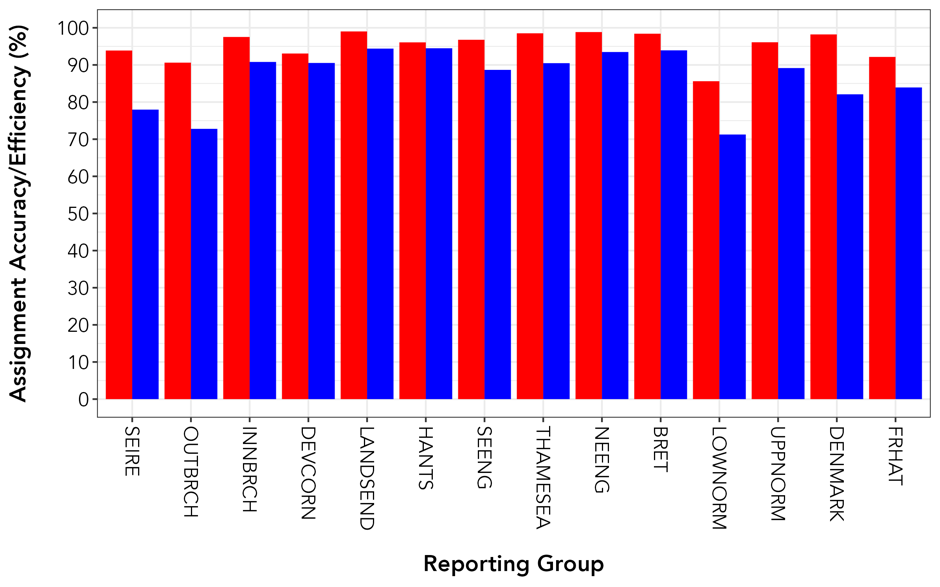

**Supplementary Figure 3** *RUBIAS* river-level Leave-One-Out accuracy (red) and efficiency (blue). River codes are as given in Table X. Reporting group codes are SEIRE – south east Ireland, OUTBRCH – outer Bristol Channel, INNBRCH – inner Bristol Channel, DEVCORN – Devon & Cornwall, LANDSEND – Land’s End, HANTS – Hampshire Basin, SEENG – south east England, THAMESEA – Thames & East Anglia, NEENG – north east England, BRET – Bretagne, LOWNORM – Lower Normandie, UPPNORM – Upper Normandie, DENMARK – Denmark and FRHAT – French hatcheries. For clarity, the Devon & Cornwall rivers have been splitting into two groups.

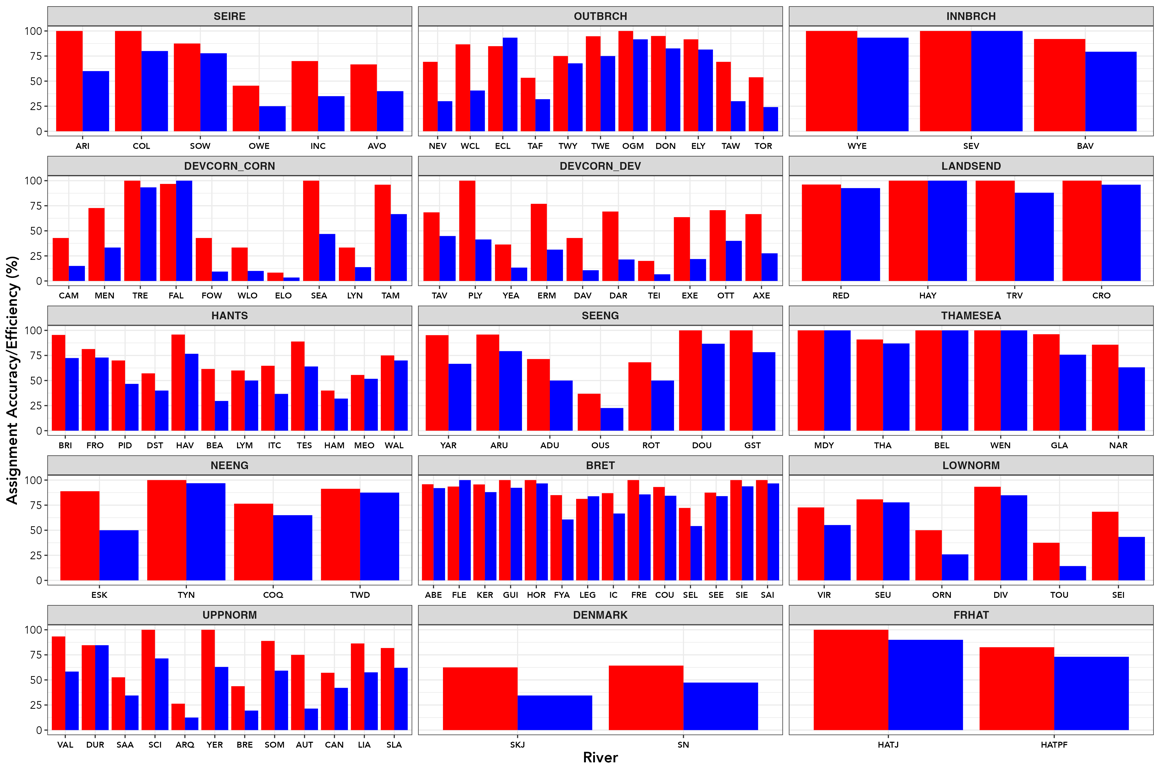

**Supplementary Figure 4** Comparison of mixed stock (MSA) and individual assignment (IA) analyses as performed in cBayes (red) and *RUBIAS* (blue) for known-origin samples of trout from 25 baseline rivers to reporting group and river of origin. River codes are as given in Supplementary Table 1. Assignment proportions are presented for the MSA and proportion of correct assignments, assuming an assignment probability ≥ 0.7 for IA.

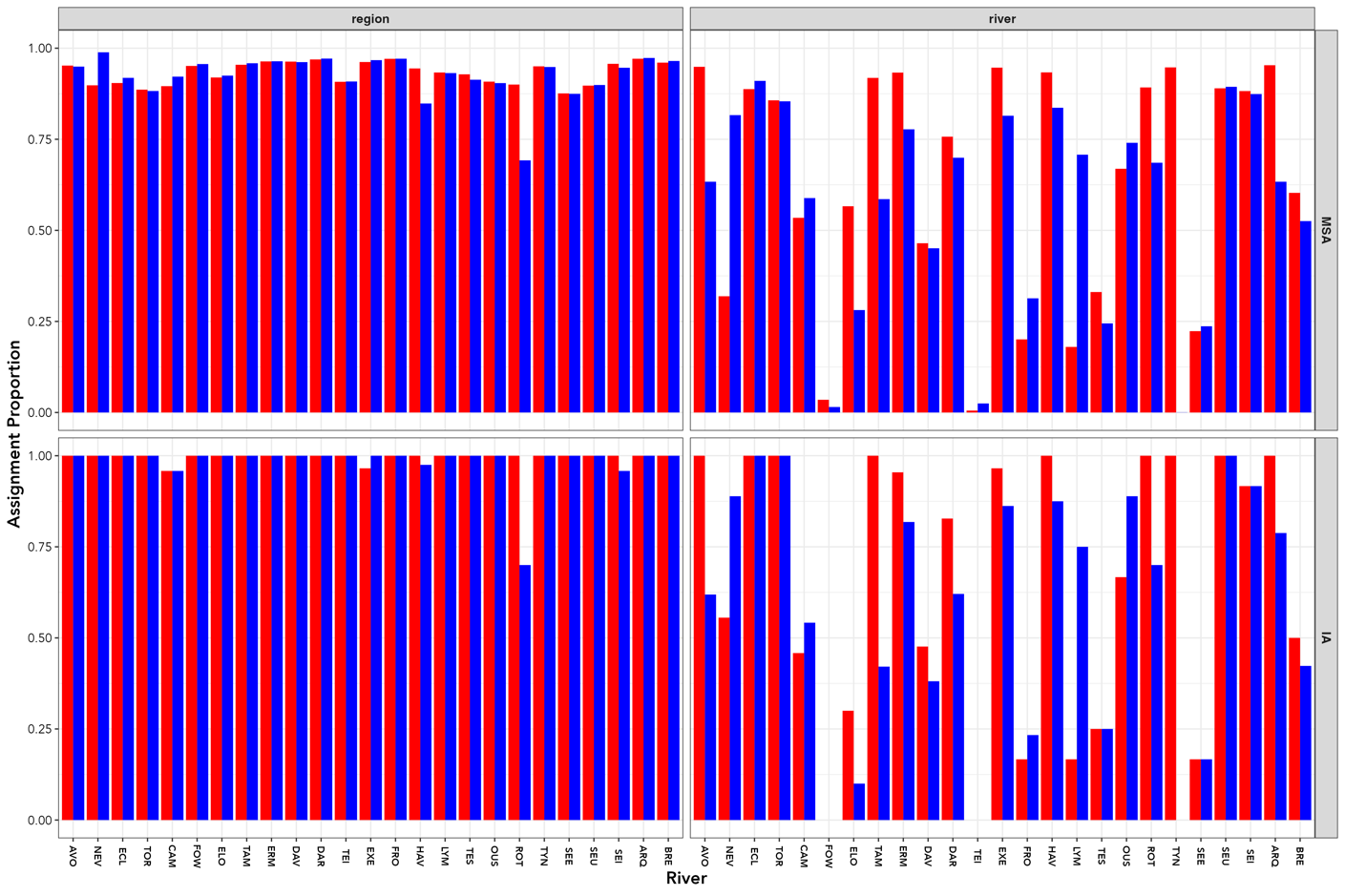

**Supplementary Figure 5** Results for Evanno et al. (2005) delta *K* (Δ*K* ) for the hierarchical STRUCTURE analyses of genetic structuring in English Channel brown trout populations.

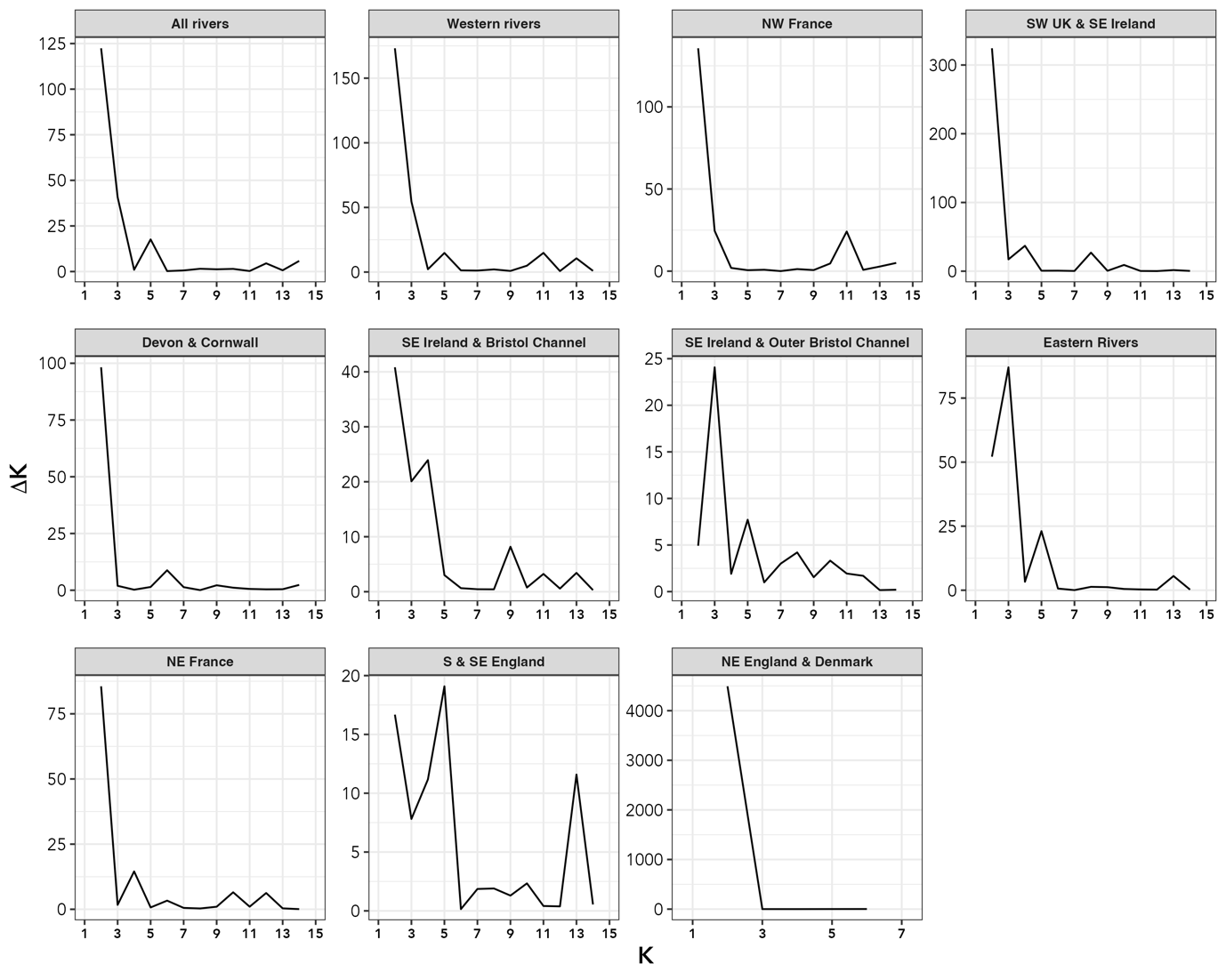
